# Energy-Saving Pretreatments Affect Pelagic *Sargassum* Composition and DNA Metabarcoding Analysis Reveals the Microbial Community Involved in Methane Yield

**DOI:** 10.1101/2023.03.21.533673

**Authors:** Enrique Salgado-Hernández, Alejandro Alvarado-Lassman, Sergio Martínez-Hernández, Jesús B. Velázquez-Fernández, Ana E. Dorantes-Acosta, Erik S. Rosas-Mendoza, Ángel I. Ortiz-Ceballos

## Abstract

*Sargassum* spp. floods the Caribbean coastlines, causing damage to the local economy and environment. These macroalgae have a low methane yield that makes the anaerobic digestion (AD) process unviable, so low-cost pretreatments are required. This research investigated the efficiency of energy-saving pretreatments, such as water washing, that had not been evaluated for these species. The microbial communities involved in AD of the best and worst-performing systems were also analyzed by high-throughput sequencing. The results showed that water washing pretreatment modified the content of inorganic compounds, fibers, and C:N ratio and increased the methane yield by 38%. The bacterial phyla Bacteroidota, Firmicutes, and Thermotogota, as well as the archaea genera *Methanosarcina*, *RumEn_M2*, and *Bathyarchaeia*, dominated the microbial communities. This study is the first to show the microbial community structure involved in the AD of *Sargassum* spp. The pretreatments presented in this study may help overcome the previously reported limitations.

## 1. Introduction

Since 2011, unusually large quantities of pelagic *Sargassum* have reached the Mexican Caribbean coast (Rodríguez-Martínez et al., 2020). Two species (one with two morphological forms) of holopelagic *Sargassum* have been identified: *S. fluitans III*, *S. natans I,* and *S. natans VIII* (García-Sánchez et al., 2020). The beaching of massive quantities of *Sargassum* spp. results in the accumulation of decomposing material that stains the coastal waters a murky brown. This is damaging not only to the tourism industry but also to the long-term health of coastal ecosystems (van Tussenbroek et al., 2017). On tourist beaches, macroalgae have been removed from the beaches and sea. However, these have been disposed of in areas that are not adequately prepared to prevent leachate from leaching into the aquifer (Rodríguez-Martínez et al., 2020). Therefore, it is necessary to find a suitable use for the large volumes of *Sargassum* biomass. Researchers have explored the valorization of these macroalgae for biofuel generation and have primarily focused on the AD to produce biogas (Milledge and Harvey, 2016; Thompson et al., 2020). Biogas production from brown algae may have a better potential energy output than biohydrogen, bioethanol, and biodiesel production. Also, the methane yields data of brown algae are higher than the terrestrial biomasses from sugar crops and lignocellulosic biomasses except for organic wastes; besides, the fact that it does not need to be cultivated makes biofuel production viable (Song et al., 2015). However, there are currently few studies on AD of these invasive species and published results report different biomethane potentials (BMP) (Milledge et al., 2020; Morrison and Gray, 2017; Tapia-Tussell et al., 2018; Thompson et al., 2020). In all cases, these values represent less than 40% of the theoretical biomethane potential (TBMP).

Low methane yields have been attributed to the compositional characteristics of *Sargassum* spp. that include a high content of minerals, ash, polyphenols, and high levels of insoluble fibers (Milledge et al., 2020; Tapia-Tussell et al., 2018). These factors, together with the low carbohydrate content, which generates low C: N ratios, have been cited as the main reasons for the low methane yield of *Sargassum* spp. (Soto et al., 2015). Brown macroalgae by themselves present high content of mineral salts (Milledge et al., 2019), including mainly light metal ions, such as sodium, potassium, calcium, and magnesium (Zhang et al., 2017a). It has been reported that at high concentrations of sodium, methane yield begins decreasing (Zhang et al., 2017b). Because of this, different pretreatments to improve the solubilization of biomass and reduce inhibitory compounds, such as mineral salts, have been assessed on several species of macroalgae (Maneein et al., 2018). For pelagic *Sargassum* spp., Tapia-Tussell et al., (2018) used a fungal pretreatment and achieved a BMP of 104 mL CH_4_ g^-1^ VS. While Thompson et al., (2020) used hydrothermal pretreatments and achieved a BMP of 116 mL CH_4_ g^-1^ VS. However, the transfer of these technologies from laboratory to industrial scale is hampered by high capital investment and energy inputs that could not be offset by methane yields. Because of this, continuous efforts are currently being made to optimize pretreatments and achieve better yields at lower cost. Freshwater washing is a low-energy pretreatment method that is commonly employed in macroalgae biofuel development. Washing has been used to remove inert contaminants and salts that can hinder methane generation in high quantities (Maneein et al., 2018). In the case of these two *Sargassum* species, washing is a common method before AD. However, its effect on methane yield and chemical composition has not been evaluated.

In addition, there is little information about the influence of macroalgae composition on the microbial community of the AD process. It is known that different bacterial and archaeal consortiums shape biomass AD, which generates complex microbial communities. These microbial communities change based on the type of substrate (Ziganshin et al., 2013), and increase their diversity with the chemical complexity of the substrate (Theuerl et al., 2019). Therefore, it is necessary to know the diversity of the microbial community that drives the AD of *Sargassum* spp.

This study investigates the use of various low-energy pretreatments based on water and chemical solutions, as well as the microbial community involved in high and low methane yields. The results show an effect on methane production, substrate biodegradability, and chemical composition that had not been reported before. The effect of *Sargassum* on the diversity of bacteria and archaea is also reported. To the best of the authors’ knowledge, this is the first study to describe the structure of the microbial composition that performs *Sargassum* spp. degradation in the AD process. The objectives of this study were: (i) to improve the biomethane potential of *Sargassum* spp. by improving its compositional characteristics using different energy-saving pretreatments; and (ii) to identify the microbial community structure associated with biomethane yield.

## 2. Materials and methods

### 2.1 Biomass and inoculum collection

*Sargassum* spp. biomass was collected in Playa del Carmen (20°37’19.99’’N, 87°4’9.98’’W), Quintana Roo, Mexico. Sand and other natural contaminants attached (plastics, shells, feathers, and other algae) to the *Sargassum*, were removed manually. Samples were transported to the laboratory in a cooler at room temperature for 4 hours and stored in resealable plastic bags at -4 °C for later use.

The inoculum was obtained from a pilot-scale reactor operating at ambient temperature and fed with the liquid fraction of municipal organic solid waste (LFMOSW), located at the Instituto Tecnológico de Orizaba (Veracruz, Mexico). The inoculum was incubated at 35°C under anaerobic conditions for 30 days and LFMORS was fed as the substrate to increase the microbial population and ensure methane production. The inoculum presented pH, TS, VS, and ash contents of 7.9±0.8, 4.15 ±0.08 %, 60.62 ±9.9% ST, and 39.4 ±9.9% ST, respectively. Before biochemical methane potential testing, a sludge sample was taken for microbial analysis.

### 2.2 Pretreatments of *Sargassum* biomass

*Sargassum* samples were subjected to three different pretreatments to improve the composition of *Sargassum* biomass. The three different pretreatments are described below:

#### 2.2.1 Water washing (WW)

The wet biomass was manually washed with fresh water at a constant flow (0.920 L min^-1^) for 2 min at room temperature (25 °C) to remove sand and other contaminants and allowed to drain for 5 min.

#### 2.2.2 Soaking + Warm water washing (S+W)

The wet biomass was immersed in fresh water at a ratio of 1: 4 (w: v) at room temperature for 24 h. Followed by a second wash with warm water at 40°C for 3 min and allowed to drain for 5 min. This method was modified based on Bruhn et al., (2011) and Tabassum et al., (2017).

#### 2.2.3 Chemical soaking (CS)

Two hundred grams of wet biomass were weighed and immersed in 400 mL of 2% formaldehyde (FHD) for a period of 16 h. The solution was drained, filtered, and rinsed with distilled water. Next, 500 mL of 0.2 M HCl was added for 24 h and again the samples were rinsed with distilled water. Finally, they were allowed to drain for 3 min. This method was modified from Fertah et al., (2017). The treatment was conducted at room temperature.

Samples of *Sargassum* biomass without pretreatment (as received), referred to as untreated (UT), were used. Finally, after pretreatments, samples were dried at 60°C to ∼10% moisture and manually ground using ceramic mortar with a pestle into powder with a particle size ≤1 mm. The powdered *Sargassum* biomass was stored in resealable plastic bags at room temperature for further analysis and evaluation of BMP.

### 2.3 Biochemical methane potential (BMP) testing

The test to evaluate the BMP was established according to Holliger et al., (2021). The bioreactors consisted of 120 mL serum bottles with a working volume of 75 mL. The untreated (UT) and pretreated (WW, S+W, CS) samples were added to each bioreactor as substrate. Powdered cellulose (Sigma-Aldrich) was used as a positive control to ensure the accuracy of the tests. Inoculum-only blanks (negative control) were also prepared to measure the contribution of biogas from the inoculum. The inoculum to substrate ratio was 2: 1 according to VS. The pH values of the cultures were adjusted to 7.2 ±0.1, at the beginning of the experiment. All bottles were hermetically sealed with butyl rubber stoppers and aluminum caps. The headspace was purged with nitrogen gas, purity 99.5-100% (Praxair México S. de R.L. de C.V) for 3 min to reach anaerobic conditions. The bottles were incubated at 32°C until the daily methane production for three consecutive days was <1% of the accumulated volume. During this period, the bottles were shaken daily for 60 s. The volume of biogas was measured at regular intervals using a 5-60 mL glass syringe with a valve and luer-lock system. The biogas produced by the bioreactors was corrected with the biogas produced by the blank. The results were presented as the volume of gas (mL) at standard conditions (273 K and 1 atm) times the mass (g) of aggregated VS. The experiment was performed in triplicate (n=3).

### 2.4 Analytical methods

Biogas composition was analyzed on a gas chromatograph (Buck Scientific 310, Norwalk, USA) with a thermal conductivity detector (TCD) equipped with a 6-inch long, 0.25-inch diameter CTR-I column. Helium was used as the carrier gas at 70 psi. The column temperature was 36°C, and the detector temperature was 121°C. TS, VS, and ash of the *Sargassum* and inoculum samples were measured according to Standard Methods (APHA, 2005). The pH was measured using a Thermo Scientific™ Orion™ Versa star meter (Waltham, USA). The total phenol content (CFT) was analyzed by the Folin-Ciocalteu method using a UV-Vis spectrophotometer (Shimadzu, UV 1280, Kyoto, Japan). The analysis of carbon, hydrogen, and nitrogen content of untreated and pretreated biomass samples was analyzed using the elemental analyzer (PerkinElmer Series II CHNS/O Analyzer 2400, Waltham, USA). The total sulfur (S) was quantified by turbidimetry with gum arabic as a stabilizer, using a UV/Vis spectrophotometer (Thermo Scientific™ Spectronic 200, Waltham, USA). The oxygen content was estimated by difference. All the above analyses were performed in triplicate.

Cellulose, hemicellulose, and lignin content were determined by detergent fiber analysis [neutral detergent fiber (NDF), acid detergent fiber (ADF), acid detergent lignin (ADL)] on an ANKOM 200 fiber analyzer (ANKOM Technology, Fairport, NY, USA). The content of the major mineral salts was determined using a flame photometer, Corning 410 (Sherwood Scientific Ltd., Cambridge, UK) for Na and K, and an atomic absorption spectrometer, Varian 240FS (Agilent Technologies, Inc., Santa Clara, USA) for Ca and Mg. The above analyses were performed in duplicate.

### 2.5 Theoretical energy value and anaerobic biodegradability index

To calculate the theoretical biomethane potential (TBMP) and the empirical formula (C_n_H_a_O_b_N_c_) of pretreated and untreated *Sargassum* spp. biomass, the Buswell equation, and the Boyle equation were used (Mahmoodi et al., 2018; Mhatre et al., 2019). The Biodegradability Index (BI) was calculated as the ratio of BMP and TBMP as the percentage of TBMP achieved by the feedstock at the end of the digestion period.

The higher heating value (HHV) was calculated according to the modified Dulong formula described by Nizami et al., (2009), and the lower heating value (LHV) was converted from the HHV as described by Deng et al., (2020).

### 2.6 Microbial community analysis

#### 2.6.1 DNA extraction

Samples (1 mL) were taken from the inoculum and the WW and CS systems at the end of BMP tests, which were selected as exemplars of high and low methanogenic performance. Samples were centrifuged at 12 000 x g for 3 min to remove the liquid. DNA from the sludge was extracted using the DNeasy PowerSoil kit from QIAGEN (Hilden, Germany) according to the manufacturer’s instructions. DNA concentration, quality, and purity were estimated using agarose gel electrophoresis (1%) and with a Nanodrop^TM^ 2000 UV-Vis spectrophotometer (Thermo Scientific, Waltham, USA).

#### 2.6.2 16S rRNA gene metabarcoding and bioinformatics analysis

Once the quality of the DNA samples was assured, PCR amplification of selected regions was conducted using specific primers connected with barcodes. Hypervariable regions V4-V5 of the 16s rRNA gene for Bacteria were amplified using primer set 515F (5 ′-GTGCCAGCMGCCGCGGTAA-3 ′) and 907R (5′-CCGTCAATTCCTTTCTTTGAGTTT-3 ′). Similarly, regions V4-V5 of the 16s rRNA gene for Archaea were amplified using the primer set Arch519F (5’-CAGCCGCCGCGGGGTAA-3’) and Arch915R (5’-GTGCTCCCCCCCGCCAATTCCT-3’). PCR products with appropriate sizes were selected by 2% agarose gel electrophoresis. The same amount of PCR products from each sample was pooled, end-repaired, A-tailed, and ligated with Illumina adapters. Libraries were sequenced on the Illumina NovaSeq 6000 platform by Novogene Co (Beijing, China). After sequencing, forward and reverse reads with barcoded and primers removed were analyzed with QIIME2 software. Demultiplexed reads were denoised and assigned to amplicon sequence variants (ASVs) using the DADA2 algorithm. Taxonomic assignment was based on classifiers trained on the V4-V5 hypervariable region extracted from the SILVA 138 99% 16S sequence database. The confidence threshold for limiting taxonomic depth was set to 0.97.

### 2.7 Statistical analysis

A one-way analysis of variance (ANOVA) was performed to investigate the effect of pretreatments on biomass composition (ash content, total phenols, and minerals) and PBM at a 95% confidence interval limit. Statistical analyses were performed with the R programming language using RStudio version 1.3.1093 (RStudio Team, 2020).

## 3. Results and discussion

### 3.1 Effect of pretreatments on the composition of *Sargassum* spp

The compositional analysis along with the energy content of untreated and pretreated *Sargassum* spp. biomass is presented in Table 1. Proximate analysis on the untreated samples (as received) showed a high ash content with a value of 43.9 ±1.3% and VS content of 44.7 ±0.9%. These results generated high ash: volatile solid ratio (A: V) of 0.98, indicating a high content of inorganic material. According to Tabassum et al., (2017), high A: V ratios can have inhibitory effects on AD. Likewise, the ratio of volatile solids to total solids (VS: TS) had a value of 0.50, which represents the organic matter content in the sample. However, after pretreatments, the ash content was significantly affected (*p* ˂0.001) and represented a decrease of approximately 50%. Whereas the VS: TS ratio increased to 0.75, 0.74, and 0.76 with WW, S+W, and CS, respectively. Thus, no significant differences were found between pretreatments (*p* >0.05). The positive effect of different pretreatments on the reduction of ash content has been reported. <u>Diaz et al., (2015)</u> tested different demineralization pretreatments of *Sargassum* spp. using different chemical solutions and all successfully decreased the amount of ash. Also, the use of washing in freshwater has been successfully reported to the reduction of ash content (Milledge et al., 2018).

**Table 1.** Effect of pretreatments on compositional characteristics of *Sargassum* spp. UT = Untreated; WW = Water washing; S+W = Soaking + Warm water washing; CS = Chemical soaking.

|  | UT | WW | S+W | CS |
| --- | --- | --- | --- | --- |
| **Proximal analysis (n=3)** |  |  |  |  |
| TS (%) | 89.1 ±0.8 | 89.9 ±0.06 | 86.3 ±0.05 | 91.8 ±3.4 |
| VS (%) | 44.7 ±0.7 | 67.3 ±1.4 | 64.0 ±0.5 | 70.2 ±3.6 |
| Ash (%) | 44.2 ±0.9 | 22.5 ±1.4 | 22.2 ±0.5 | 21.6 ±1.8 |
| VS: ST ratio | 0.50 | 0.75 | 0.74 | 0.76 |
| A: V ratio* | 0.98 | 0.33 | 0.35 | 0.31 |
| **Ultimate analysis (% TS) (n=3)** |  |  |  |  |
| C | 21.9 ±2.4 | 33.4 ±0.4 | 34.6 ±1.7 | 37.6 ±1.3 |
| H | 2.2 ±0.5 | 4.9 ±0.06 | 4.9 ±0.3 | 5.6 ±0.2 |
| O | 24.5 ±3.3 | 35.0 ±0.6 | 32.3 ±2.0 | 30.3 ±1.6 |
| N | 1.7 ±0.4 | 1.5 ±0.2 | 2.0 ±0.13 | 2.8 ±0.3 |
| S | 0.33 ±0.03 | 0.29 ±0.01 | 0.31 ±0.03 | 0.32 ±0.03 |
| C: N ratio | 13.2 ±1.6 | 22.2 ±2.5 | 17.0 ±1.6 | 13.6 ±1.2 |
| **Structural composition (% TS) (n=2)** |  |  |  |  |
| Cellulose | 3.7 ±0.60 | 13.2 ±0.20 | 12.0 ±0.89 | 15.2 ±1.5 |
| Hemicellulose | 8.2 ±0.59 | 10.1 ±1.20 | 11.9 ±0.56 | 8.0 ±0.90 |
| Lignin | 7.5 ±0.85 | 10.7 ±0.60 | 11.9 ±0.90 | 28.6 ±3.2 |
| TPC** | 4.3 ±0.2 | 4.7 ±0.07 | 3.5 ±0.16 | 1.6 ±0.18 |
| **Energy content** |  |  |  |  |
| HHV (kJ·g<sup>-1</sup> VS)* | 6.2 | 12.1 | 13.0 | 15.3 |
| LHV (kJ·g<sup>-1</sup> VS)* | 5.5 | 10.8 | 11.7 | 13.9 |
| TBMP (mL CH<sub>4</sub>·g<sup>-1</sup> VS) * | 335.8 | 424.8 | 452.7 | 503.8 |
| Chemical formula | C<sub>1.8</sub>H<sub>2.15</sub>O<sub>1.53</sub>N<sub>0.12</sub> | C<sub>2.8</sub>H<sub>4.9</sub>O<sub>2.18</sub>N<sub>0.10</sub> | C<sub>2.9</sub>H<sub>4.9</sub>O<sub>2.02</sub>N<sub>0.15</sub> | C<sub>3.13</sub>H<sub>5.6</sub>O<sub>1.9</sub>N<sub>0.19</sub> |

\*A: V = Ratio of ash to volatile solids; HHV = Higher heating value; LHV = Lower heating value; TBMP = Theoretical biomethane potential.
\*\*TPC = Total phenol content expressed as gallic acid equivalents (GAE) per gram of dry weight. Data are mean ± SD.

The C: N ratio of the untreated samples (13:1) was lower than that registered by Milledge et al., (2020) and Thompson et al., (2020) for *Sargassum* spp. A high C: N ratio indicates the accumulation of carbohydrates, which could be easily degraded during AD while a low C: N ratio indicates a high protein content, resulting in excessive ammonia production, which causes toxicity to the methanogenic community (Milledge et al., 2019). After pretreatments, WW managed to improve the C: N ratio, reaching a value of 22:1 followed by SW with a value of 17:1. While chemical soaking (CS) generated a low C: N ratio of 13:1, similar to that of the UT samples. According to these results, only WW pretreatment would be suitable for AD since the optimum C: N ratio for microbial growth is known to be 20-30:1. However, it is reported that the optimum C: N ratio can vary with algal species, from 14:1 to 30:1 (Milledge et al., 2020).

The amount of cellulose, hemicellulose, and lignin for UT biomass was 3.7 ±0.6, 8.2 ±0.6, and 7.5 ±0.8%, respectively. These results showed that about 40% of the organic compounds in the algal biomass are composed of fibers. It is known that brown macroalgae are characterized by a low lignin content and a high amount of carbohydrates, which makes them an attractive feedstock for AD (Amador-Castro et al., 2021). However, the presence of lignin-like materials and lignified tissue in *Sargassum* spp. has been reported (Alzate-Gaviria et al., 2021). In previous works, Salgado-Hernández et al., (2023) reported a lignin content of 16.8% while <u>Tapia-Tussell et al., (2018)</u> reported a value of 15.6%. Fig. 1 shows the components of the fiber content as part of the organic matter compared to the inorganic components after pretreatments. It is possible to observe how the content of cellulose increased three times with WW and SW, due to ash removal. Hemicellulose and lignin did not show a change after pretreatments. However, CS pretreatment increased up to four times the fiber content compared to UT. Mainly, the lignin content more than doubled compared to the other pretreatments. This result can be attributed not only to the removal of inorganic compounds but also to the reduction of some soluble carbohydrates.

**Fig. 1.**
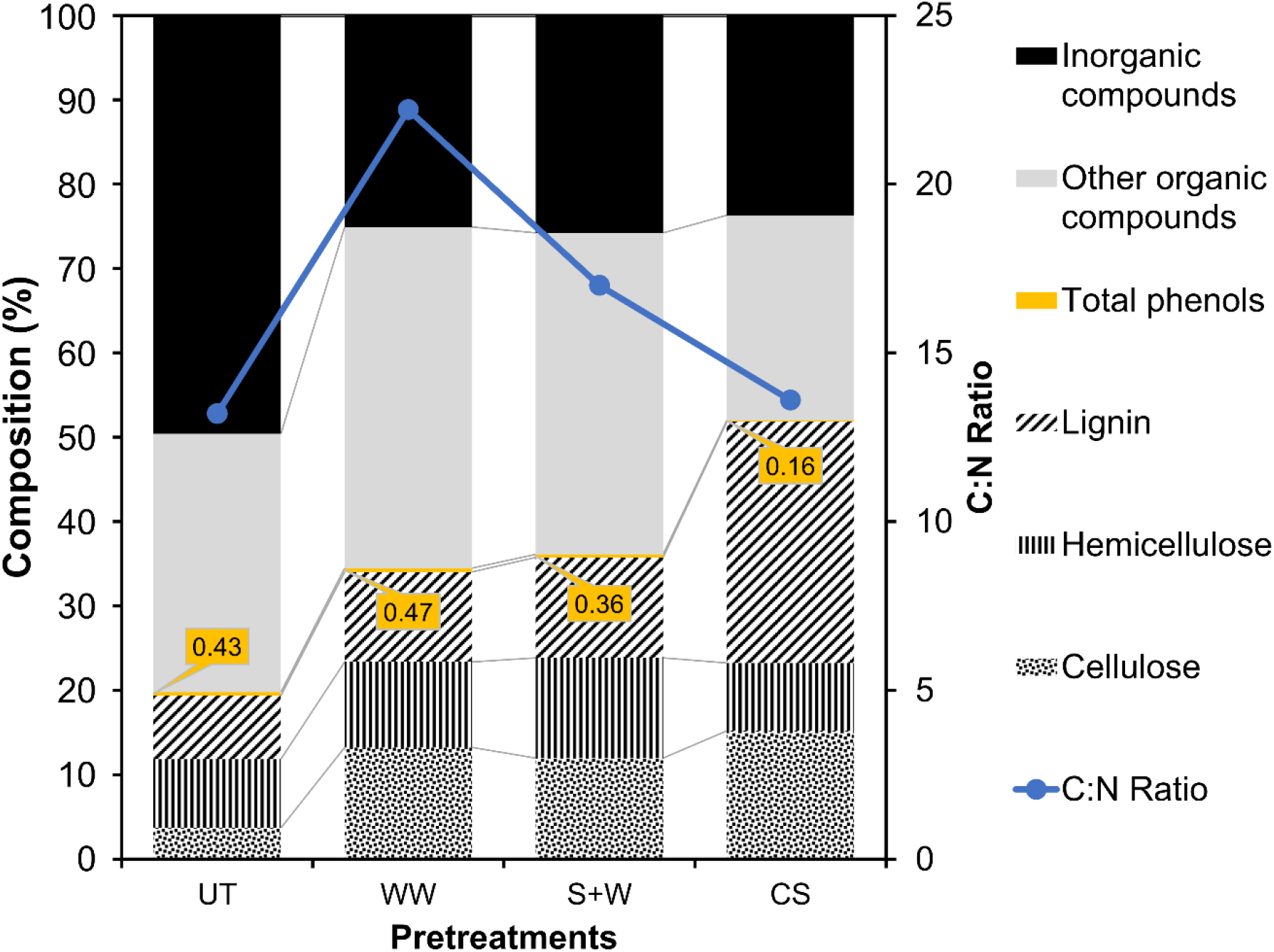
Changes in the compositional characteristics (% dry weight) of *Sargassum* spp. by the effect of pretreatments. UT = Untreated, WW = Water washing, S+W = Soaking + Warm water washing, CS = Chemical soaking.

The total phenol content (TPC) was 4.3 ±0.27 mg AGE·g^-1^ dry weight for the untreated samples. The results obtained in this study are close to those reported by Milledge et al., (2020) and Davis et al., (2021) for *Sargassum* spp. After pretreatment, TPC showed significant differences (*p* =0.000158). The SW and CS pretreatments achieved a reduction of 19 and 64%, respectively. The result obtained using CS can be attributed to the use of formaldehyde, which can bind phenolic compounds (Moen et al., 1997). Only in the case of WW pretreatment, the difference from UT biomass was not significant (*p* =0.2794076).

### 3.2 Effect of pretreatments on mineral content

Table 2 shows the results of mineral content, mainly light metal ions of Na, K, Ca, and Mg, present in untreated and pretreated *Sargassum* biomass. The UT samples presented a high content of Ca, followed by Na, K, and to a lesser extent Mg. This result is similar to that reported by Davis et al., (2021) and Rodríguez-Martínez et al., (2020) who also found a high content of Ca, followed by K, Na, and Mg. The total mineral content in this study was 125.4 mg·g^-1^, with Ca contributing 60% of the total content. With the application of the pretreatments, a significant reduction in the total content of these minerals was achieved (*p* =0.00000372), as shown in Table 2. The CS pretreatment achieved the most significant reduction of minerals (Na, K, Ca, and Mg) compared to UT biomass (*p* =0.0000002), which represented a removal of 57%. <u>Ross et al., (2009)</u> also reported removals of Mg, K, Na, and Ca in brown algae of up to 90% with the use of an acid pretreatment, while water pretreatment achieved reductions of 30-40%. The pretreatment S+W results better than the pretreatment WW for reducing Na and K content, which is advantageous as they have been reported to govern the toxicity of methanogens (Milledge et al., 2019). However, WW and S+W did not present significant differences between them, as both achieve a similar reduction of minerals.

**Table 2.** Influence of pretreatments on the content of mineral salts and total removal in the biomass of *Sargassum* spp. UT = Untreated, WW = Water washing, S+W = Soaking + Warm water washing, CS = Chemical soaking.

|  | Na | K | Ca | Mg | Total* | Removal |
| --- | --- | --- | --- | --- | --- | --- |
|  | mg·g<sup>-1</sup> dry weight |  |  |  |  | % |
| **UT** | 32 ±0.2 | 10.9 ±1.2 | 74.8 ±2.5 | 7.7 ±0.9 | 125.4 |  |
| **WW** | 23.4 ±0.9 | 7.9 ±0.1 | 48.1 ±1.9 | 8.9 ±0.2 | 88.3 | 30 |
| **S+W** | 6.6 ±0.5 | 2 ±0.09 | 70.8 ±1.3 | 10.8 ±0.3 | 90.2 | 28 |
| **CS** | 4.1 ±0.06 | 0.4 ±0.1 | 46.9 ±1 | 2.8 ±0.13 | 54.2 | 57 |

Data represent mean ± SD, n=2.
\*The data sets were analyzed with a one-way ANOVA showing a $p=0.00000372$ .

### 3.3 Effect of pretreatments on the theoretical energy content of *Sargassum* spp

The TBMP of the untreated samples was 335.8 mL CH_4_·g^-1^ VS and after pretreatment, the TBMP improved up to 50%, as shown in Table 1. The increase of TBMP appears to be associated with reduction of ash and minerals in algal biomass. Table 1 shows an increase in the percentages of organic elemental composition, mainly C and H, after pretreatments. This change in elemental composition modifies the stoichiometric formula and thus the amount of methane that can be predicted. Mainly, the CS pretreatment was the one that reached the highest value, possibly due to the removal of inorganic matter with the use of chemical solutions. Hessami et al., (2019) and Mhatre et al., (2019) also found an increase in TBMP after the removal of some bioproducts using different acidic and alkaline solutions. On the other hand, the HHV of biomass doubled after pretreatments, with the CS pretreatment achieving a value 2.4 times higher than that of the untreated *Sargassum* biomass. These results are attributed to the removal of inorganic material, such as ash content (Milledge et al., 2014).

### 3.4 Effect of pretreatments on the biomethane potential of *Sargassum* spp

The average values obtained for the accumulated biogas, methane, and biodegradability index for each pretreatment are shown in Table 3. After 53 days, the BMP of the cellulose (CL) used as a positive control produced 374.4 ±7.5 mL CH_4_·g^-1^ VS, with which we can validate the results since it fits the criteria established by Holliger et al., (2021).

**Table 3.** A summary of the results of the biomethane potential test of untreated and pretreated *Sargassum* spp. Biogas and methane accumulated after 70 days of incubation and 53 days for cellulose.

| Substrate | Biogas Yield<br>(N mL·g<sup>-1</sup><br>VS) | Methane<br>content<br>(%) | Methane<br>yield*<br>(N mL·g<sup>-1</sup><br>VS) | BI<br>(%) | Pretreatment<br>efficiency (%) |
| --- | --- | --- | --- | --- | --- |
| CL | 578.6 ±18.9 | 55.7 | 374.52 ±7.5 | 90.2** | - |
| UT | 131.53 ±3.7 | 61.7 | 77.64 ±3.9 | 23.09 | - |
| WW | 193.19 ±15.6 | 56.0 | 107.42 ±6.7 | 32.11 | +38 |
| S+W | 176.27 ±7.5 | 54.9 | 97.75 ±5.4 | 29.08 | +26 |
| CS | 150.4 ±5.4 | 49.1 | 74.77 ±5.4 | 22.23 | -4 |

CL = Cellulose, UT = Untreated, WW = Water washing, S+W = Soaking + Warm water washing, CS = Chemical soaking.
\* $p$ -value =0.000121.
\*\*BI according to the TBMP of cellulose (Filer et al., 2019).

In contrast to the control, all experiments stabilized and reached the final methane production, at 70 days of incubation. Net cumulative methane production can be observed in Fig. 2. The methane production of the UT biomass was 77.64 ±3.9 mL CH_4_·g^-1^ VS. The pretreatments had a significant effect on BMP (*p* =0.000121). The best result was obtained with WW pretreatment, which generated a BMP of 107.42 ±6.7 mL CH_4_·g^-1^ VS, which represented an increase of 38% compared to the UT biomass. SW pretreatment had a result close to WW, which represented an increase of 26%. However, the BMP between WW and S+W pretreatment were statistically similar (*p* =0.1706440). These results can be attributed to the reduction in the inorganic matter content and the increase in the C: N ratio because of the pretreatments. Yields similar to that achieved in this study were reported by Tapia-Tussell et al., (2018) and Thompson et al., (2020). In contrast, CS pretreatment, despite presenting the best reduction of inorganic matter and total phenol content in the biomass, negatively affected the BMP. CS generated the lowest BMP, which represented a decrease of 4%. CS did not show significant differences compared to UT (*p* =0.8891660). This result may be due to a possible reduction of soluble carbohydrates, which was reflected in the increase in fiber content, mainly lignin, and in a low C: N ratio, which can be seen in Fig. 1. In this study the phenol content had no effect on BMP, possibly because the phenol content was lower than that reported by other authors (Milledge et al., 2020; Tapia-Tussell et al., 2018).

**Fig. 2.**
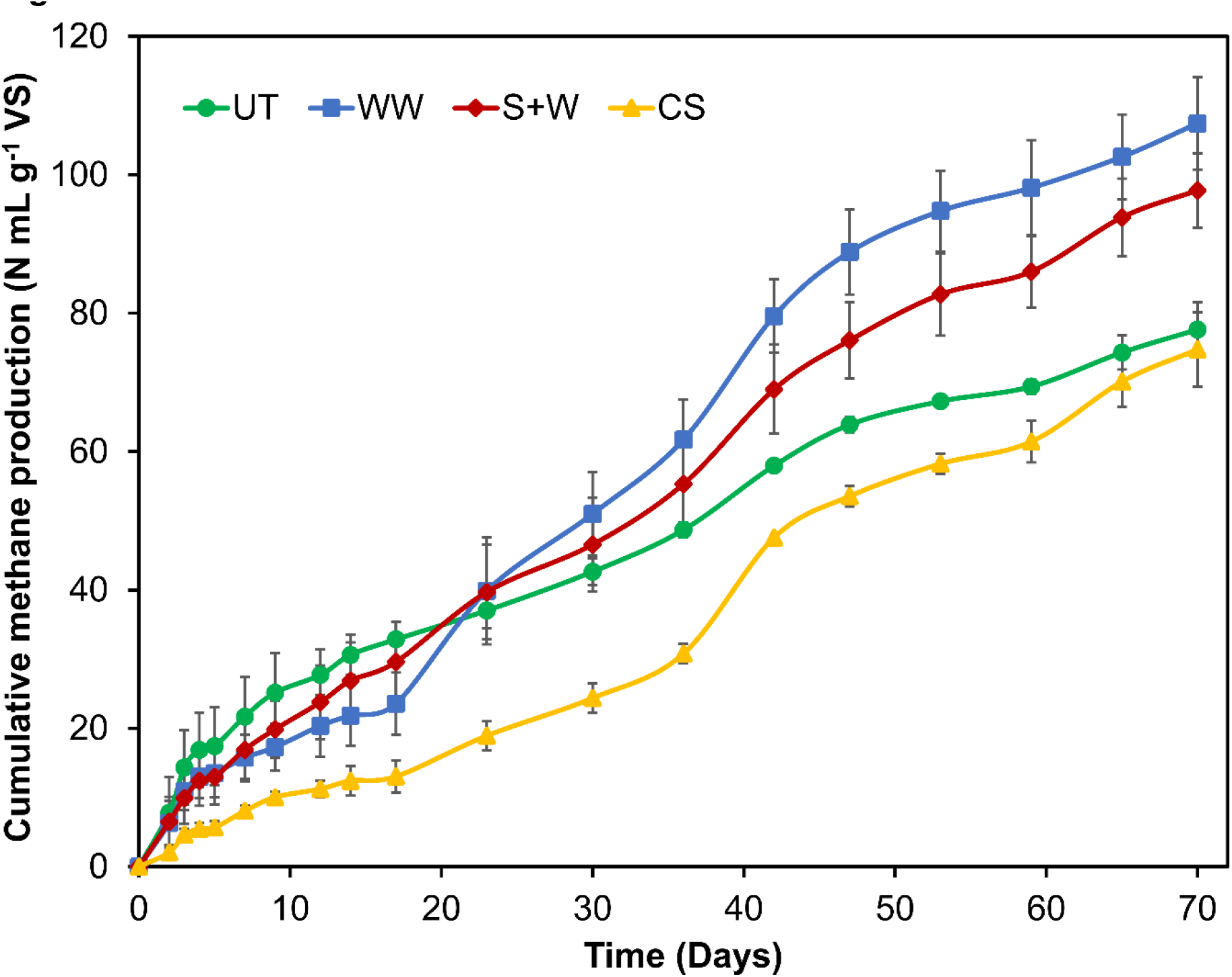
Cumulative methane production expressed in N mL·g^-1^ VS of pretreated and untreated *Sargassum* spp. biomass after 70 days of incubation. UT =Untreated, WW =Water washing, S+W = Soaking + Warm water washing, CS=Chemical soaking. Error bars represent the standard deviation of the mean (n = 3).

WW and S+W pretreatments succeed only in boosting methane yield; however, they did not improve the digestion rate, as the duration of the PBM assay was longer than that observed in previous studies (Milledge et al., 2020; Tapia-Tussell et al., 2018; Thompson et al., 2020). Furthermore, when observing the methane production curves (Fig. 2), they present a staggered shape over time, indicating that the hydrolysis of the material is the limiting step (Filer et al., 2019). The above was reflected in the biodegradability index, since the values achieved were less than 50% (Table 3). This low biodegradability index can be attributed to the recalcitrance of some organic polymers and the presence of a significant amount of fiber in the *Sargassum* spp. Therefore, studies focused on improving the hydrolysis of *Sargassum* spp. biomass is needed.

Although the methane yields achieved in this study are similar to those reported by other authors, our results were achieved using energy-saving methods. This technology requires a lower energy input than other physical pretreatment methods and eliminates the need for expensive chemicals and enzymes. Thus, this method is advantageous for addressing the problem of massive flooding of pelagic *Sargassum* affecting Caribbean beaches. One potential disadvantage of WW pretreatment is the use of freshwater since this is a natural resource and its responsible use should be considered.

### 3.5 Microbial community analysis

The analysis of the microbial community structure was conducted on the WW and CS reactors and the inoculum. Alpha-diversity indices that reflect diversity (Shannon and Simpson indices) and abundance (Chao 1) are shown in Table 4. The high numerical values of bacterial sequences and ecological indices indicated that bacterial diversity and richness greatly exceeded that of archaea.

**Table 4.**
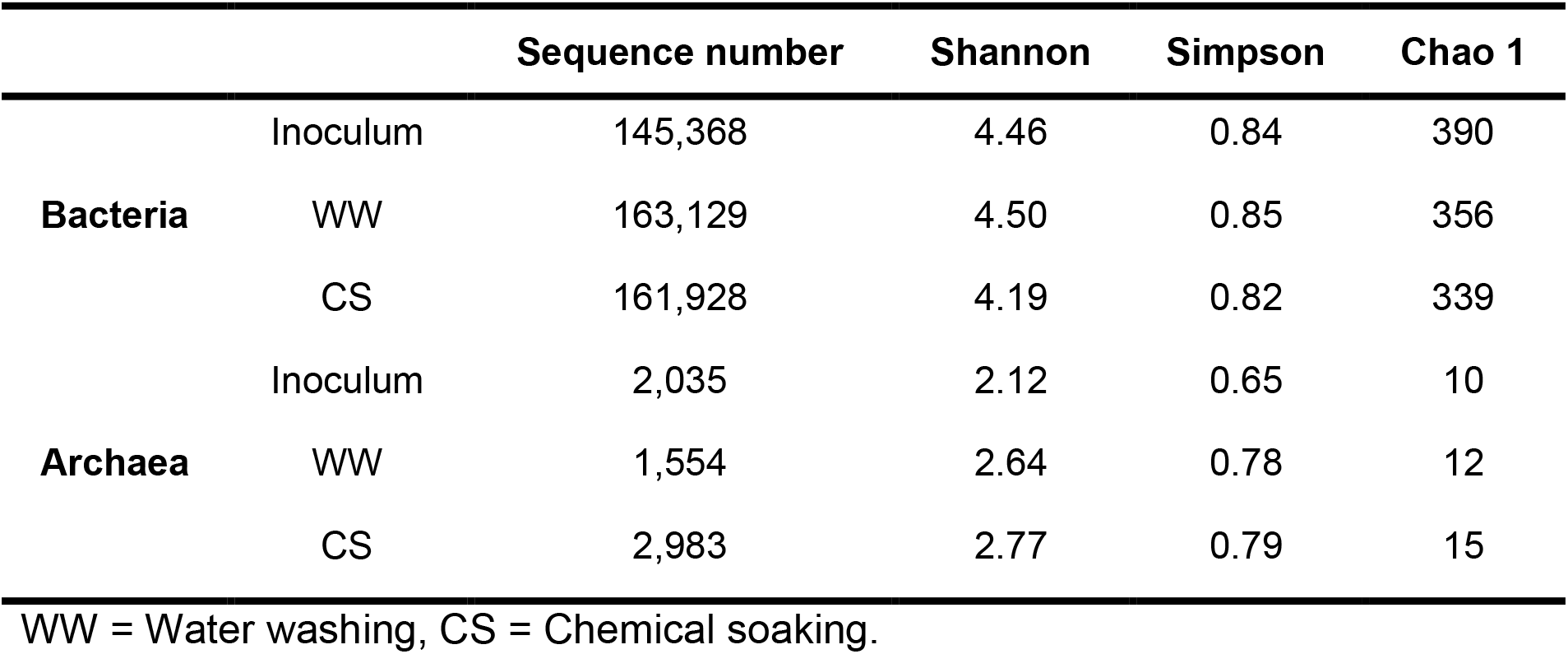
Alpha-diversity indices of the different microbial communities.

The relative abundances in the bacterial communities at the phylum and genus level are shown in Fig. 3. In both the inoculum and WW and CS reactors, no differences in dominant taxa were observed at the phylum level. All three systems were dominated by the phyla Bacteroidota (64-69%), Firmicutes (12-15%), Thermotogota (3-9%), Planctomycetota (2-4%), Synergistota (2-3%), Spirochaetota (2-5%), and Chloroflexi (1-2%).The microorganisms assigned to these taxa convert complex macromolecular organic compounds into biodegradable molecules (Amha, 2018). Therefore, these taxa are usually present in the stages of hydrolysis and acidogenesis. The phylum Firmicutes also represents the syntrophic bacteria, which are involved in the acetogenic phase (Amha, 2018). Studies show that the presence of members of the phylum Bacteroidota is dominant in systems with readily biodegradable feedstocks (Theuerl et al., 2019). However, the biomass of *Sargassum* spp. is characterized by the low availability of biodegradable compounds. Therefore, the dominance of the phylum Bacteroidota in the WW and CS systems is attributed to the inoculum source, which was fed with the liquid fraction of RSOM, which is characterized by high biodegradability.

**Fig. 3.**
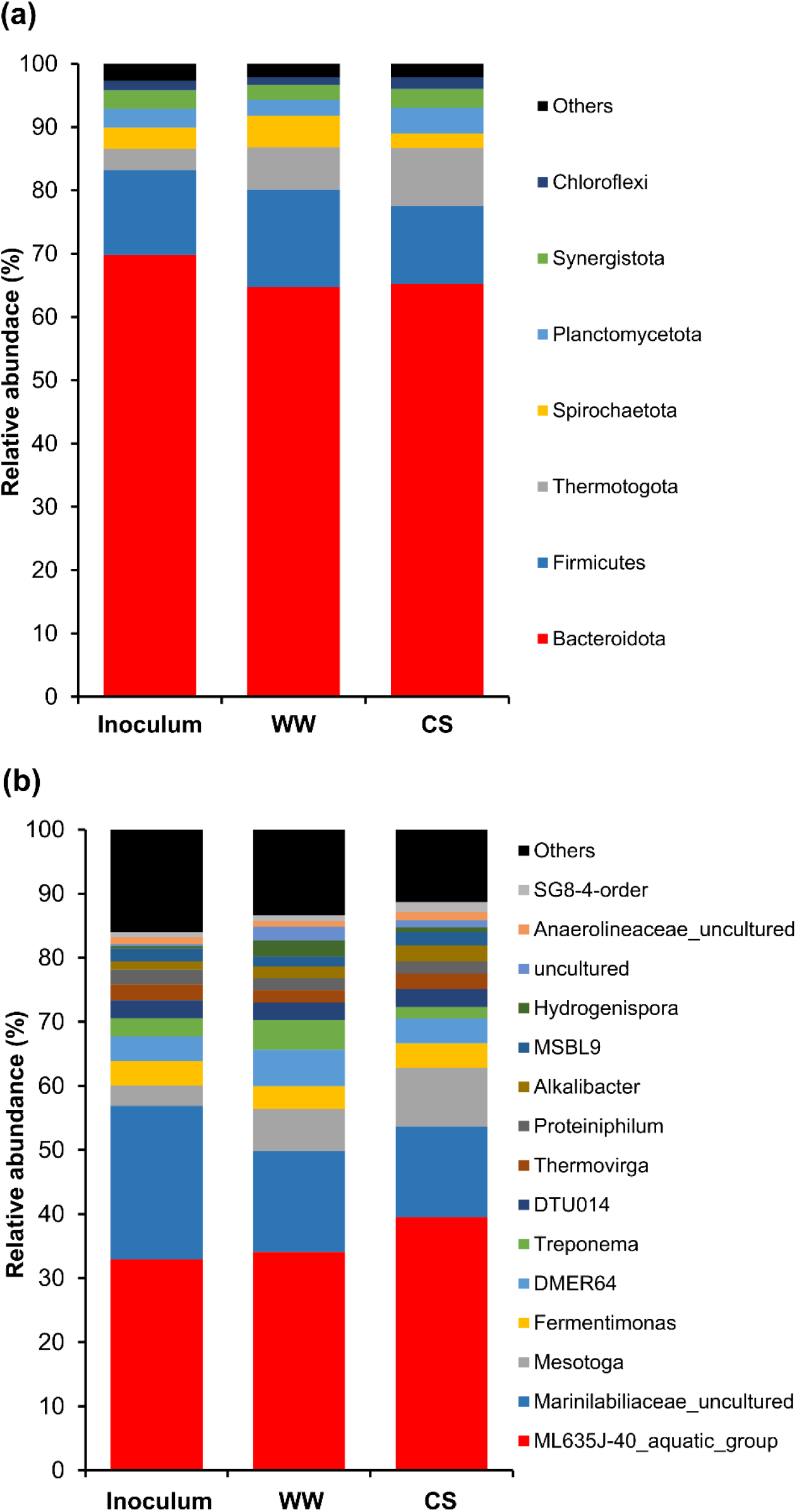
Relative abundances of bacterial communities at the phylum (a) and genus (b) level of the inoculum, WW (water washing), and CS (chemical soaking). Taxa with abundances <1% of the total sequences in the samples were grouped as other.

In Fig. 3(a), at the genus level, *ML635J-40_aquatic_group*, dominated in both the inoculum (32%) and the WW (34%) and CS (39%) systems. This group of microorganisms was assigned to the ML635J-40_aquatic_group family that is characterized by conducting the fermentation of organic materials such as sugars, starches, or more complex polysaccharides (Wan et al., 2018). Another group of bacteria, classified as *Marinilabiliaceae_uncultured*, belonging to the Marinilabiliaceae family was abundant in the inoculum (24%). However, a decrease was observed in the WW (16%) and CS (14%) systems. Members of this family are characterized by being saccharolytic, i.e., they ferment sugars (Whitman et al., 2015). Therefore, we can attribute the decrease in Marinilabiliaceae to the scarcity of fermentable sugars in the CS and WW media. In contrast, an increase in the abundance of the genus *Mesotoga* was observed in WW (7%) and CS (9%) compared to the inoculum (3%). This genus is abundant in mesothermal anaerobic environments rich in aromatic compounds. It can perform H_2_ oxidation and thiosulfate reduction using elemental sulfur as a final electron acceptor (Nesbø et al., 2019). Therefore, the slight increase in *Mesotoga* can be attributed to the presence of aromatic compounds derived from the decomposition of lignocellulosic material in WW and CS.

The results of the analysis of the archaeal community structure at the phylum and genus level are shown in Fig. 4. It was found that most of the 16S rRNA gene reads from both the inoculum and WW and CS were assigned to the phylum Holobacterota (47-65%), Thermoplasmatota (28-37%), and Crenarchaeota (6-13%). Currently, Halobacterota and Thermoplasmatota contain most of the known methanogens.

**Fig. 4.**
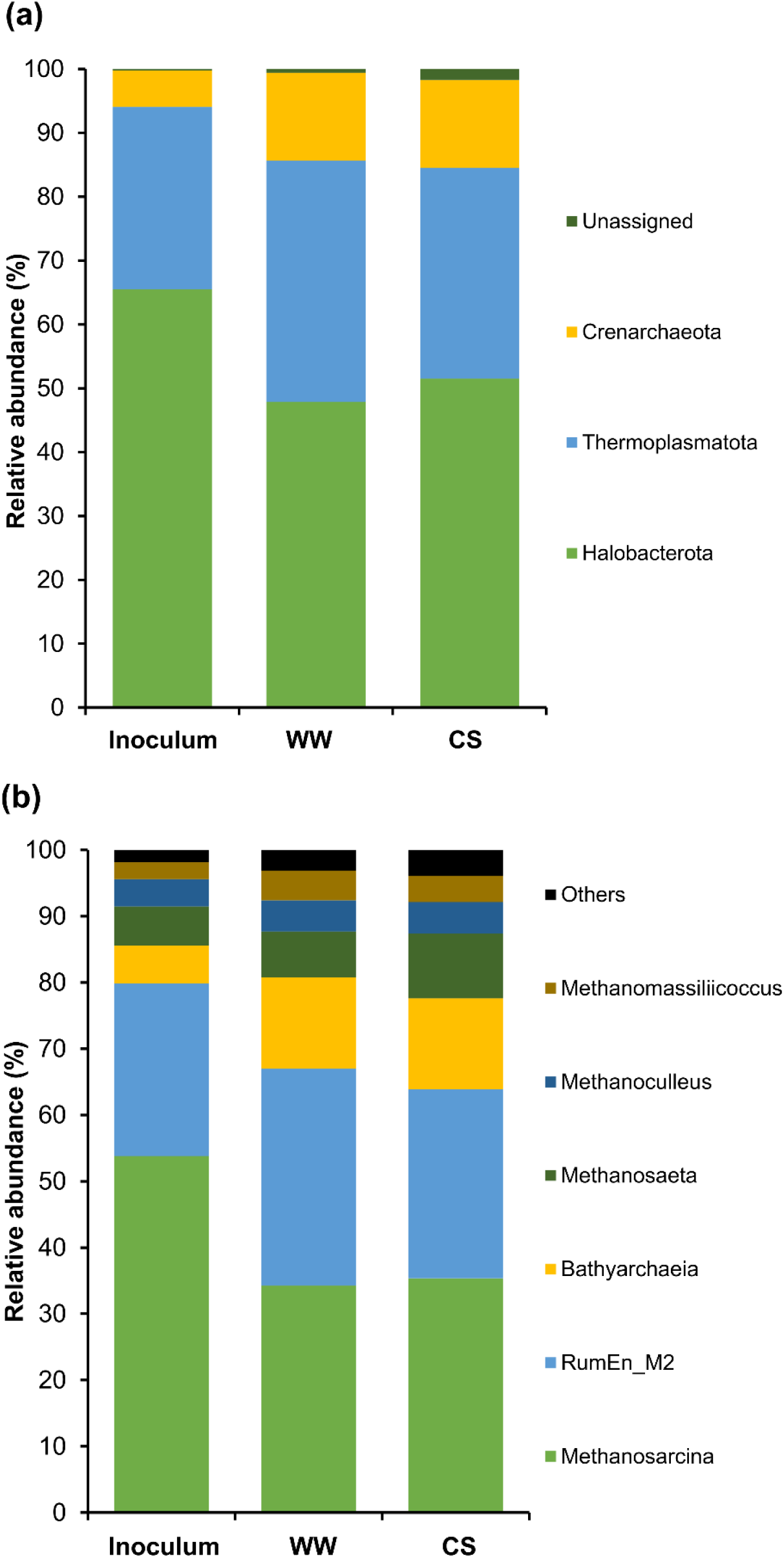
Relative abundances of archaeal communities at the phylum (a) and genus (b) level of the inoculum, WW (water washing), and CS (chemical soaking). Taxa with abundances <2% of the total sequences in the samples were grouped as other.

At the genus level, *Methanosarcina*, *RumEn_M2*, *Bathyarchaeia*, *Methanosaeta*, *Methanoculleus*, and *Methanomassiliicoccus* were detected (Fig. 4b). These genera are known to play an important role in methanogenesis, except *RumEn_M2* and *Bathyarchaeia* for which until a few years ago there was little information on their involvement in methanogenesis. The genus *RumEn_M2* (family methanomethylophilaceae), is now known to be a new methanogenic genus, which is restricted to growth on methanol and methylamines and is strictly H_2_-dependent (Jiang et al., 2022). Also, the presence of putative methane-metabolizing genes in members of Bathyarchaeia suggests that methanogenesis may be more widespread phylogenetically than we know (Vanwonterghem et al., 2016).

*Methanosarcina*, which are mixotrophic methanogens and can metabolize acetate, hydrogen, and C_1_ compounds, were dominant in the inoculum (54%). However, the relative abundance of *Methanosarcina* decreased in WW (34%) and CS (35%) systems, despite being characterized by high tolerance to stress conditions and environmental changes. This decrease can be attributed to the low acetogenic activity in WW and CS media. High concentrations of acetate are known to favor the growth of *Methanosarcina* (Wang et al., 2018). Therefore, there was a slight increase in the genus *Methanosaeta* in WW (7%) and CS (10%) compared to the inoculum (6%), which is usually more abundant at low acetate concentrations due to its high affinity with the substrate. However, an increase in the relative abundance of *Bathyarchaeia* from 6% (inoculum) to 14% was observed in WW and CS. Li et al., (2021) reported that *Bathyarchaeia* can perform the same functions as some hydrogenotrophic methanogens and therefore displace them due to their high affinity for H_2_ and CO_2_. That ability of *Bathyarchaeia* could reduce the abundance of *Methanosarcina*, which was dominant in the inoculum but decreased in abundance in WW and CS after *Bathyarchaeia* increased. These results showed that *Sargassum* spp. decreased the activity of the acetoclastic pathway and increased the hydrogenotrophic pathway. Possibly due to the low number of fermentable compounds by the hydrolytic bacteria that reduced the activity of the acidogenic and acetogenic bacteria, due to the low biodegradability of *Sargassum* spp.

The microbial composition present in the inoculum source remained the same during AD of *Sargassum* spp. only the presence of *Sargassum* spp. favored the growth of the genera *Mesotoga* and *Bathyarchaeia*. Since the WW and CS systems presented a similar microbial composition, it was impossible to identify the microorganisms responsible for high and low methane production. It was observed that methane production from *Sargassum* spp. was more dependent on the substrate composition, as was the high amount of insoluble fibers and low C: N ratio in the CS system.

## 4. Conclusions

Pretreatments succeeded in modifying the composition of the *Sargassum* spp. WW had a positive effect on inorganic matter content, C: N ratio, and fiber content, which were key to increasing methane yields. CS pretreatment was the best in the removal of mineral salts and in the improvement of the theoretical energy content, however, it did not have an improvement in the experimental methane yield. Although it is important to reduce the inorganic compounds in *Sargassum* spp. biomass, the content of insoluble fibers continues to be a barrier in biodegradability and methane production. Therefore, pretreatments focused on improved hydrolysis of this lignocellulosic biomass are required.

The microbial community composition involved in AD of *Sargassum* spp. appears to depend on the inoculum source. No differences were found in the composition of the microbial communities of the systems with high and low methanogenic yields. The pretreated *Sargassum* spp. favored the growth of *Mesotoga* and *Bathyarchaeia*, displacing the genus *Methanosarcina*. This study also suggests the use of a different inoculum source with greater affinity to the algal biomass.

## Acknowledgments

The authors thank the federal maritime-terrestrial zone (ZOFEMAT) of the municipality of Solidaridad (Quintana Roo, México) for obtaining the macroalgae. Enrique Salgado Hernández thanks CONACYT for a doctoral scholarship (817679).

## Notes

### Competing Interest Statement

The authors have declared no competing interest.

